# Sennoside A and Ceftazidime Inhibit Nucleocapsid RNA Binding Across SARS-CoV-2, SARS-CoV, and MERS-CoV

**DOI:** 10.64898/2026.06.09.731089

**Authors:** Shweta Singh, Gagan D Gupta

## Abstract

SARS-CoV, MERS-CoV, and SARS-CoV-2 exemplify the persistent threat posed by coronaviruses, with their capacity for zoonotic spill over, rapid transmission, and high mortality, and thus underscores the urgent need for broad-spectrum antiviral strategies. The nucleocapsid (N) protein, essential for RNA binding, genome packaging, and viral replication, is highly conserved among coronaviruses but remains an underexplored antiviral target. In our earlier work, we identified two small molecules, ceftazidime and sennoside A, that bind the N-terminal domain of the SARS-CoV-2 N protein and inhibit nucleic acid binding, and identified their binding sites using NMR chemical shift perturbation assays. Here, we observed that several residues involved in inhibitor binding are conserved across betacoronaviruses, suggesting a shared druggable vulnerability. We have purified recombinant N proteins from SARS-CoV, MERS-CoV, and SARS-CoV-2, and demonstrated by electrophoretic mobility shift assays that both compounds significantly reduced RNA binding. Their inhibitory concentrations (*IC*_*50*_) were determined using fluorescence polarization. The docking analyses indicated that both inhibitors target the RNA-binding pocket of the N-NTD, consistent with a conserved mechanism of action. Collectively, our findings reveal a conserved RNA-binding vulnerability in coronavirus N proteins and highlights the pan-coronavirus therapeutic potential of these inhibitors.

## Introduction

Coronaviruses pose a persistent and significant threat to global health due to their capacity for zoonotic transmission, rapid adaptation, and the ability to cause severe respiratory illnesses in humans [1–3]. Over the past two decades, three highly pathogenic betacoronaviruses have emerged: Severe Acute Respiratory Syndrome coronavirus (SARS-CoV) in 2002, Middle East Respiratory Syndrome coronavirus (MERS-CoV) in 2012, and Severe Acute Respiratory Syndrome coronavirus 2 (SARS-CoV-2) in 2019 [1,4,5]. While SARS-CoV and MERS-CoV were responsible for localized but severe outbreaks, SARS-CoV-2 led to the COVID-19 pandemic, resulting in unprecedented global morbidity, mortality, and socio-economic disruption [6– 8]. These recurring outbreaks underscore the urgent need for broad-spectrum antiviral strategies and sustained pandemic preparedness [8–10].

Coronaviruses possess large, positive-sense single-stranded RNA genomes of approximately 30 kb, which encode both non-structural and structural proteins [11,12]. The non-structural proteins, including the RNA-dependent RNA polymerase (nsp12), main protease (nsp5), and helicase (nsp13), are essential for viral replication and have been the primary focus of antiviral drug development [11–13]. The structural proteins— spike (S), envelope (E), membrane (M), and nucleocapsid (N)—play critical roles in viral entry, assembly, and genome packaging. The S protein is particularly important for host entry: SARS-CoV and SARS-CoV-2 utilize angiotensin-converting enzyme 2 (ACE2) as their receptor, whereas MERS-CoV engages dipeptidyl peptidase 4 (DPP4/CD26) [12,13]. Despite differences in receptor usage and sequence divergence, these coronaviruses share conserved genome organization and essential replication machinery, underscoring common vulnerabilities that can be exploited for therapeutic development [14].

Historically, antiviral strategies have focused on targeting viral components prone to rapid mutation, such as the spike protein or essential viral enzymes like the RNA-dependent RNA polymerase (RdRp) and main protease (Mpro) [14]. These approaches have led to the development of therapeutics including remdesivir, molnupiravir, and neutralizing monoclonal antibodies [15–17]. However, the long-term efficacy of these interventions is challenged by the emergence of viral variants with mutations that confer drug resistance or immune escape, as well as concerns regarding off-target toxicity and inconsistent clinical performance across different patient populations [16–19].

In contrast, the Nucleocapsid (N) protein—a multifunctional RNA-binding protein essential for genome packaging, replication, and immune modulation—offers a more conserved target among coronaviruses [20–23]. The N protein is among the most abundantly expressed viral proteins and demonstrates high sequence and structural preservation, especially within its N-terminal (NTD) and C-terminal (CTD) domains [24]. Moreover, it forms phase-separated condensates facilitating viral assembly and antagonizes host antiviral stress responses, making it central to viral pathogenesis [22,23,25].Structural studies have revealed that the N-terminal domain (NTD) of the N protein contains a conserved RNA-binding groove across betacoronaviruses, characterized by a β-sheet “palm” and a basic β-hairpin “finger,” which are critical for selective recognition of viral RNA elements [24,26,27].Despite its essential functions and high degree of conservation, the N protein remains an underexplored antiviral target.

Our previous work has identified two FDA-approved small molecules, ceftazidime and sennoside A, that bind to the SARS-CoV-2 N-NTD and disrupt its interaction with RNA [28]. These inhibitors primarily engage conserved residues within the β-hairpin loop and palm region, as determined through NMR chemical shift perturbation experiments [28]. These observations raise the possibility of broad-spectrum activity of these inhibitors against other human coronaviruses. Building on these findings, the present study investigates the conservation of the N-NTD RNA-binding pocket across SARS-CoV-2, SARS-CoV, and MERS-CoV, and evaluates the ability of ceftazidime and sennoside A to inhibit RNA binding by N proteins from all three viruses. By integrating sequence analysis, biochemical assays, and molecular docking, we demonstrate that these inhibitors target a conserved and druggable vulnerability in the coronavirus N protein, supporting the development of pan-coronavirus therapeutics.

## 2. Materials and methods

### 2.1. Generation of partial domain constructs of N protein

To enable comparative functional studies of conserved RNA-binding domains, we generated partial domain constructs of the nucleocapsid (N) protein from SARS-CoV-2, SARS-CoV, and MERS-CoV. The SARS-CoV-2 construct, encoding residues 44–364 (SCoV2-N44–364) and encompassing both the N-terminal domain (NTD) and C-terminal domain (CTD) while excluding disordered terminal regions, was cloned into the pNH-TrxT expression vector (Addgene plasmid #26106) as previously described [28]. This construct enables expression of recombinant SCoV2-N44–364 with an N-terminal 6xHis tag, a thioredoxin (Trx) fusion partner, and a TEV protease cleavage site for tag removal following purification.

Similarly, codon-optimized gene fragments encoding the N proteins from MERS-CoV (MCoV-N_39–362_; residues 39–362, UniProt ID: A0A1W5LGD9) and SARS-CoV (SCoV-N_49–365_; residues 49–365, UniProt ID: P59595) were chemically synthesized and cloned into the lab-engineered vector pPCS-Trx-TEV expression vector having pET-28a(+) backbone, and modified to include an N-terminal 6×His tag, Trx fusion partner, and TEV protease site. The final construct sequences are provided in the supplementary materials. This cloning strategy ensured consistency in downstream protein purification and assay conditions. All constructs were verified by Sanger sequencing and subsequently used for recombinant protein expression, purification, and functional characterization in assays such as RNA binding and inhibitor interaction studies.

### 2.2. Expression and purification of Nucleocapsid proteins

The nearly full-length nucleocapsid (N) protein constructs— SCoV-N_49–365_, SCoV2-N_44–364_, and MCoV-N_39–362_ were heterologously expressed in *E. coli* Rosetta™ cells (Merck Life Science, India). The cells were cultured in Lysogeny Broth (LB) medium supplemented with kanamycin (50 μg/mL) and chloramphenicol (25 μg/mL) at 37 °C until reaching an optical density at 600 nm (OD_600_) of 0.8. Protein expression was induced by adding 0.5 mM isopropyl β-D-1-thiogalactopyranoside (IPTG), followed by overnight incubation at 18 °C to enhance soluble expression. The cells were harvested through centrifugation, and the cell pellets were re-suspended in ice-cold lysis buffer [25 mM Tris-HCl pH 8.0, 500 mM NaCl, 25 mM imidazole, 10 % glycerol, 0.5 mg/mL lysozyme, 1 mM PMSF, and a protease inhibitor cocktail (cOmplete tablets, Roche)]. Cell disruption was achieved via ultrasonication on ice, and the lysate was clarified by centrifugation at 15,000 x g for 45 min at 4 °C. All proteins, containing an N-terminal 6xHis tag, were purified individually using an immobilized metal chelating affinity column (Ni-IDA, GE Healthcare) pre-charged with 200 mM NiSO4 and equilibrated with buffer H1 [25 mM Tris-HCl pH 8.0, 500 mM NaCl, 25 mM imidazole]. The proteins were eluted using a linear gradient of 25–500 mM imidazole over 6 column volumes using a Biologic LP system (BioRad). The additional tag residues (6xHis-Thioredoxin tag) were cleaved using in-house purified TEV Protease in a buffer solution [25 mM Tris-HCl pH 8.0, 100 mM NaCl] by incubating overnight at 4 °C. The resulting protein samples were subsequently loaded onto a Ni-IDA affinity column, allowing the pure protein, devoid of any additional residues, to be collected as the unbound fraction. The proteins were further purified using gel-filtration chromatography using Superdex™ 200 column (GE Healthcare) as the final purification step. The protein fractions were analysed using 12 % SDS-PAGE. The proteins were concentrated using ultra-centrifugal filters with 30 kDa cut-offs (Millipore). Protein concentrations were determined using a Nano Photometer (Implen) based on the absorbance at 280 nm and the molar extinction coefficients determined using the ExPASy ProtParam tool (http://www.expasy.org).

### 2.3. Size exclusion chromatography

The purified N protein constructs were analyzed by size-exclusion chromatography (SEC) on a Superdex™ 200 10/300 GL column (GE Healthcare) pre-equilibrated with SEC buffer (25 mM sodium phosphate, pH 7.0; 100 mM NaCl). The column was calibrated using standard molecular weight markers: alcohol dehydrogenase (150 kDa), bovine serum albumin (66 kDa), carbonic anhydrase (29 kDa), and cytochrome C (12.4 kDa). Fractions corresponding to the major single peak were collected and subsequently used for biophysical and biochemical analyses.

### 2.4. Dynamic light scattering

The hydrodynamic size and homogeneity of recombinant N-protein constructs (SCoV-N_49–365_, SCoV2-N_44–364_, and MCoV-N_39–362_) were assessed by dynamic light scattering (DLS) using a Zetasizer Nano-ZS (Malvern Instruments, UK). Proteins concentrations were adjusted to 1 mg/mL into SEC buffer, and centrifuged at 20,000 x *g* for 10 min at 4 °C. DLS measurements were performed at 25 °C with three runs per sample, each comprising 12 consecutive 10s acquisitions. Intensity-weighted size distributions and polydispersity indices (PDI) were calculated using the Zetasizer software, and mean hydrodynamic diameters (Dh) represent averages of the three runs.

### 2.5. Nucleic acid binding assays

The RNA-binding activity of the N proteins from SARS-CoV-2, SARS-CoV, and MERS-CoV was evaluated using electrophoretic mobility shift assay (EMSA). A 15-mer RNA oligonucleotide (5′-CGGUUUCGUCCGUGU-3′), corresponding to a region within the SARS-CoV-2 untranslated region (UTR), was used as the RNA probe. The RNA was generated via in vitro transcription and purified before use. For binding assays, increasing concentrations of recombinant N protein constructs, SCoV-N_49–365_, SCoV2-N_44–364_, and MCoV-N_39–362_ were incubated with 100 ng of RNA probe in a 10 μL reaction volume in SEC buffer. The mixtures were incubated at 4 °C for 30 minutes to allow protein– RNA complex formation, and resolved on 1.5% agarose gel pre-stained with ethidium bromide. Gels were imaged using a UV transilluminator to detect shifts in RNA bands indicating protein–RNA interactions.

To assess the inhibitory potential of small molecules on RNA binding, EMSAs were also performed in the presence of candidate inhibitors—ceftazidime, and sennoside A. For inhibition assays, 10 μM of purified N protein was pre-incubated with increasing concentrations of each inhibitor (ranging from 50 to 800 μM) in the reaction buffer on ice for 30 minutes. Subsequently, 100 ng of RNA probe was added and the reaction was allowed to proceed for an additional 30 minutes. Samples were then resolved on agarose gels under identical conditions. A reduction in RNA–protein complex formation in the presence of inhibitors indicated their efficacy in disrupting the RNA-binding function of the N protein.

### 2.6. Determination of RNA Binding Affinity (KD) and Inhibitory Potency (IC50) of Nucleocapsid Protein Constructs

Fluorescence anisotropy (FA) was employed to determine the equilibrium dissociation constants (*K*_*D*_) of N protein constructs: SCoV-N_49–365_, SCoV2-N_44–364_, and MCoV-N_39–362_, using the protocol described earlier [29].A 15-mer FAM-labelled single-stranded RNA probe (5′-FAM-CGGUUUCGUCCGUGU-3′) was used at a final concentration of 20 nM. Protein-RNA binding reactions were performed in SEC buffer in 10µl volume, with increasing concentrations of each unlabelled N protein (0–5000 nM). Reactions were incubated for 30 minutes at 4 °C to allow equilibrium binding. Fluorescence polarization (FP) measurements were recorded using a CLARIOstar® microplate reader (BMG Labtech) at excitation and emission wavelengths of 485 nm and 525 nm, respectively. Each experiment was performed in triplicate, and averaged values were used for analysis. The resulting FP values were converted to anisotropy and plotted against protein concentration. Binding curves were fitted using a one-site specific binding model in GraphPad Prism (version 8.4.2) to calculate the apparent *K*_*D*_ values.

For inhibitor studies, a fixed protein concentration of 2000 nM—corresponding to near-saturation binding conditions—was selected. N protein constructs, SCoV-N_49–365_, SCoV2-N_44–364_, and MCoV-N_39–362_ were pre-incubated with increasing concentrations (0–400 μM) of ceftazidime and sennoside A, in the same SEC buffer for 30 minutes at 4 °C. After inhibitor binding, 20 nM FAM-labelled RNA probe was added to each reaction, followed by incubation for another 30 minutes at 4 °C. FP signals were recorded as described above. Anisotropy values were plotted against the logarithm of inhibitor concentrations, and the half-maximal inhibitory concentrations (*IC*_*50*_) were determined by fitting the data to a four-parameter sigmoidal dose-response model using GraphPad Prism. This fluorescence anisotropy-based approach enabled precise quantification of the RNA-binding affinity of N protein homologs and assessment of small-molecule inhibitors targeting RNA interaction.

### 2.7. Molecular Docking of Inhibitors to the Nucleocapsid N-Terminal Domain

To predict the binding modes of ceftazidime and sennoside A targeting the RNA-binding site of the N-terminal domain (NTD) of the coronavirus nucleocapsid (N) protein, molecular docking simulations were performed using the HADDOCK 2.4 web server [30,31]. A single protomer from the available crystal structures of the N-NTD of SARS-CoV-2 (PDBID 8X1H), SARS-CoV (PDBID 1SSK), and MERS-CoV (PDBID 4UD1) were selected for docking. All water molecules were removed prior to docking to optimize binding surface accessibility. The 3D structures of the ceftazidime and sennoside A were obtained from the ZINC database and LigandBox, respectively [32,33]. Among the key residues involved in inhibitor interaction identified from NMR chemical shift perturbation studies of SARS-CoV-2 N-NTD, only those conserved across the N-NTDs of SARS-CoV and MERS-CoV were designated as active residues in the docking protocol to maintain consistency for cross-species docking analysis [28]. The top-ranked docking poses were further evaluated for residue-level interactions and overall binding energy. Binding affinities of the resulting protein–ligand complexes were predicted using the PRODIGY web server (https://wenmr.science.uu.nl/prodigy/), which estimates free energy of binding (ΔG) based on interfacial contacts [34].

## 3. Results

### 3.1. The Coronavirus Nucleocapsid Protein Exhibits High Sequence Conservation Across Pathogenic Strains

To assess the evolutionary conservation of the nucleocapsid (N) protein among highly pathogenic human coronaviruses, we performed multiple sequence alignments (MSA) of full-length N proteins from SARS-CoV-2, SARS-CoV, and MERS-CoV, using clustal omega (https://www.ebi.ac.uk/jdispatcher/msa/clustalo) (**Fig. 1**) [35].

**Figure 1.**
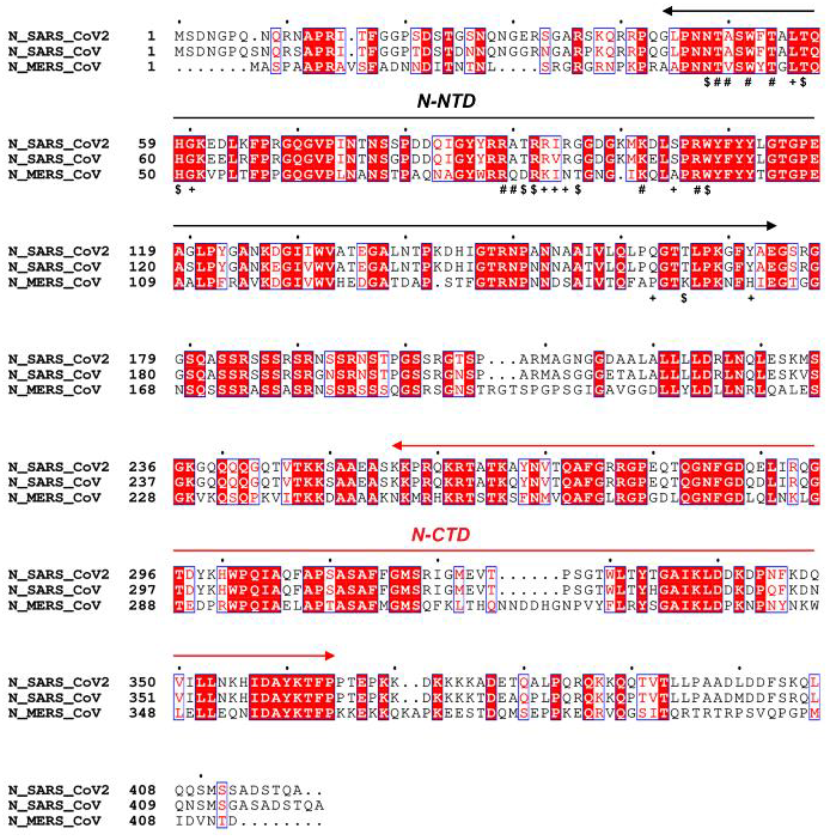
Multiple sequence alignment of nucleocapsid proteins from SARS-CoV-2, SARS-CoV, and MERS-CoV. Identical residues are highlighted, while positions containing similar residues are boxed. The N-NTD and N-CTD domains are indicated above the alignment.

SARS-CoV-2 N shares approximately 90 % sequence identity with SARS-CoV and 49 % with MERS-CoV, underscoring its critical and conserved role in viral genome packaging and replication. The N-terminal domain (NTD), the RNA binding domain, of SARS-CoV-2 shares ∼92 % identity with SARS-CoV N-NTD, and ∼59% identity with MERS-CoV N-NTD (**Fig. 1**). Further we mapped the residues perturbed by inhibitors binding, identified previously by NMR chemical shift perturbation (CSP) for SARS-CoV-2 N-NTD, onto the alignment (**Fig. 1**) [28]. These residues—including L56, H59, G60, R92, R93, I94, W108, and A173 for ceftazidime, and N48, T49, A50, W52, T54, T57, H59, R89, R92, K102, R107, W108, and E118 for sennoside A—are highly conserved across all three viruses. This conservation suggests that inhibitors targeting SARS-CoV-2 N-NTD may also be effective against other human coronaviruses.

### 3.2. Expression, Purification, and Biophysical Characterization of Nucleocapsid Proteins

Recombinant N protein constructs— SCoV-N_49–365_, SCoV2-N_44–364_, and MCoV-N_39–362_— were expressed in *E. coli* and purified to near homogeneity (**Fig. 2A**). Following tag removal, analytical size-exclusion chromatography confirmed that all constructs exist as homodimers in solution, with estimated molecular weights of ∼70 kDa, consistent with previous reports [12,24,28].

**Figure 2.**
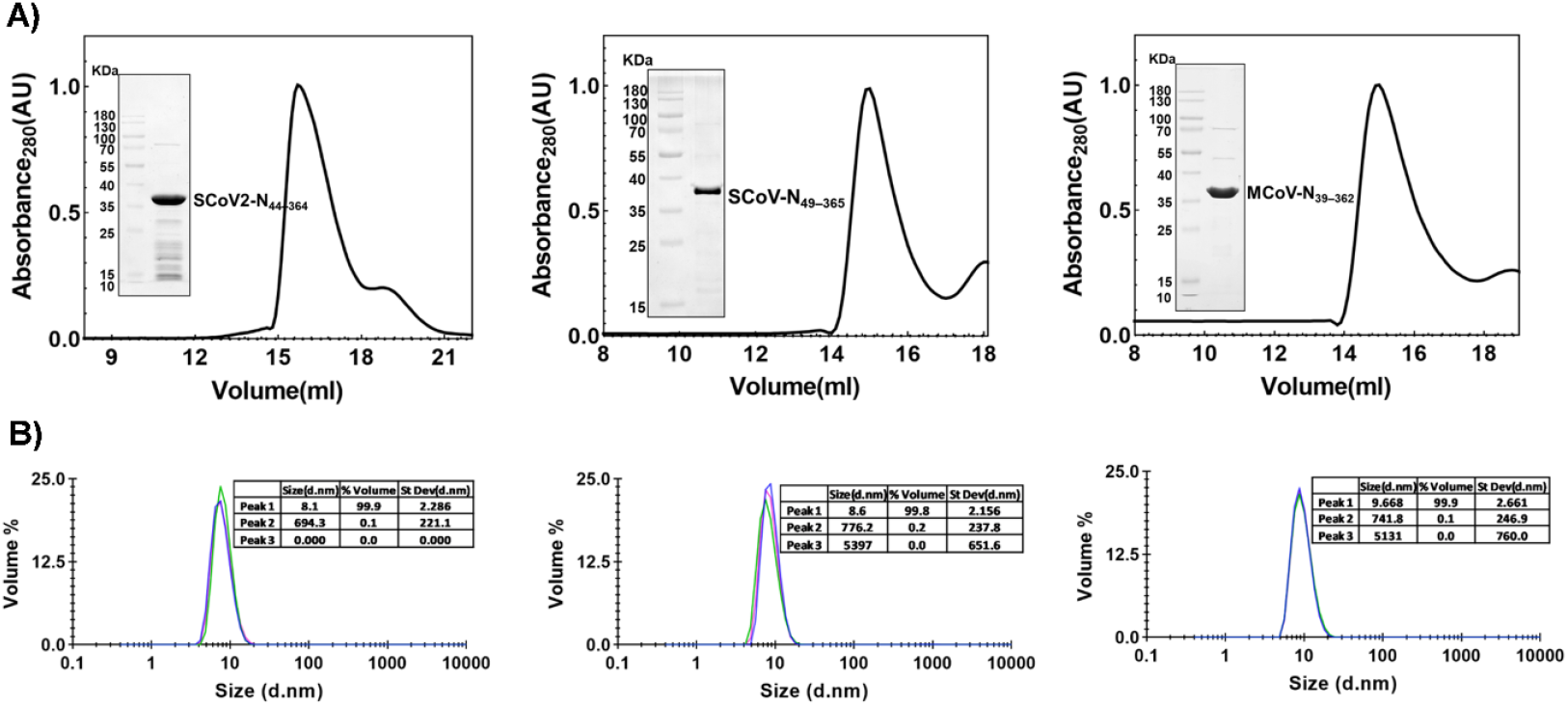
Purification and characterization of all three Nucleocapsid proteins. A) Size-exclusion chromatography (SEC) profiles and corresponding SDS–PAGE analyses (insets) of purified N protein constructs—SARS-CoV-2 N_44–364_, SARS-CoV N_49–365_, and MERS-CoV N_39–36_—comprising the structured NTD, CTD, and IDR2 but excluding the flanking IDRs. Each construct exhibits a single, symmetric elution peak consistent with a dimeric assembly. B) Dynamic light scattering (DLS) analysis confirms the homogeneity of the constructs, revealing a dominant monodisperse population (>99%) for each protein.

Dynamic light scattering analysis revealed a single major species (>99% intensity) for each construct, with hydrodynamic radii of 8.6 ± 2.1 nm (SCoV-N_49–365_), 8.1 ± 2.2 nm (SCoV2-N_44–364_), and 9.6 ± 2.6 nm (MCoV-N_39–362_) (**Fig. 2B**). The moderate polydispersity indices (PDI 0.3–0.4) likely reflect the intrinsic flexibility of the central disordered linker region.

### 3.3. All Three Nucleocapsid Proteins Bind RNA with High Affinity

The RNA-binding activity of the N protein constructs was evaluated using EMSA with a 15-mer RNA probe derived from the5′ UTR region of SARS-CoV-2 genome, known to contain a conserved transcriptional regulatory sequence [29]. All three constructs displayed robust RNA binding, as evidenced by retarded migration of the RNA probe at protein concentrations as low as 2.5 μM. Complete shift to slower-migrating RNA–protein complexes were observed at ≥10 μM, indicating tight binding and possible oligomerization (**Fig. 3A–C**).

**Figure 3.**
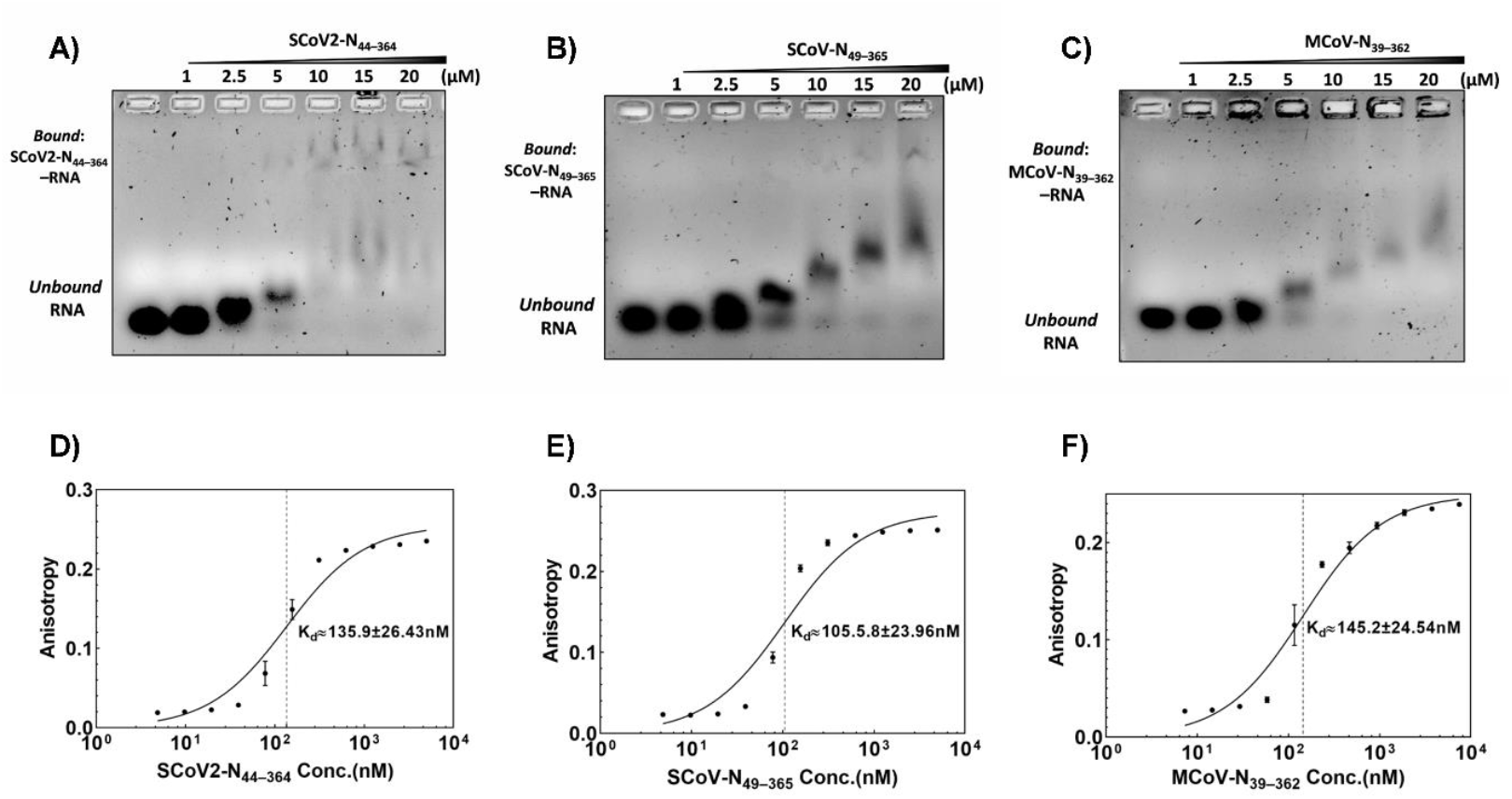
RNA-binding activity of nucleocapsid protein constructs from SARS-CoV-2, SARS-CoV, and MERS-CoV. (A– C) Electrophoretic mobility shift assays (EMSAs) showing concentration-dependent binding of SCoV2-N_44–364_, SCoV-N_49–365_, and MCoV-N_39–362_ to a 15-mer SARS-CoV-2 5′UTR RNA probe, evidenced by progressive formation of slower-migrating RNA–protein complexes. (D–F) Equilibrium binding analysis by fluorescence anisotropy using a 5′-FAM-labeled 15-mer RNA probe with the corresponding purified recombinant nucleocapsid protein constructs.

We further evaluated the RNA binding affinity of the all three N proteins using fluorescence polarization, as reported earlier [29]. Increasing concentrations of each N construct were titrated against a fixed amount (20 nM) of 5′-FAM-labeled SARS-CoV-2 UTR RNA probe (15-mer), and the assays further confirmed high-affinity RNA binding. The dissociation constants (*K*_*D*_) for SCoV2-N_44–364_, SCoV-N_49–365_ and MCoV-N_39–362_ were 135.9 ± 26.4 nM, 105.6 ± 24.0 nM, and 145.2 ± 24.5 nM, respectively (**Fig. 3D–F, Table 1**). Since saturation of RNA binding was observed at approximately 2 μM protein concentration for all nucleocapsid homologs, this concentration was subsequently used in the FP-based inhibition assays.

**Table 1.** Quantitative analysis of RNA binding and inhibitor potency for nucleocapsid protein constructs.

| S. N | N protein constructs | RNA-Binding Affinity $K_D$ (nM) | Ceftazidime ($IC_{50}$ in $\mu\text{M}$) | Sennoside A ($IC_{50}$ in $\mu\text{M}$) |
| --- | --- | --- | --- | --- |
| 1 | SCoV2-N44–364 | 135.9±26.43 | 48.49 ± 2.83 | 44.71 ± 5.34 |
| 2 | SCoV-N49–365 | 105.58±23.96 | 71.13 ± 2.05 | 57.95 ± 3.33 |
| 3 | MCoV-N39–362 | 145.2±24.54 | 77.96 ± 7.46 | 56.58 ± 13.74 |

### 3.4. Sennoside A and Ceftazidime Inhibit RNA Binding of Nucleocapsid Homologs

We next assessed whether sennoside A and ceftazidime could disrupt RNA binding by the N proteins. EMSA experiments demonstrated dose-dependent inhibition for both compounds. Ceftazidime effectively inhibited SARS-CoV-2 N–RNA binding at at 200 μM with near-complete inhibition at 400 μM (**Fig. 4A**).

**Figure 4.**
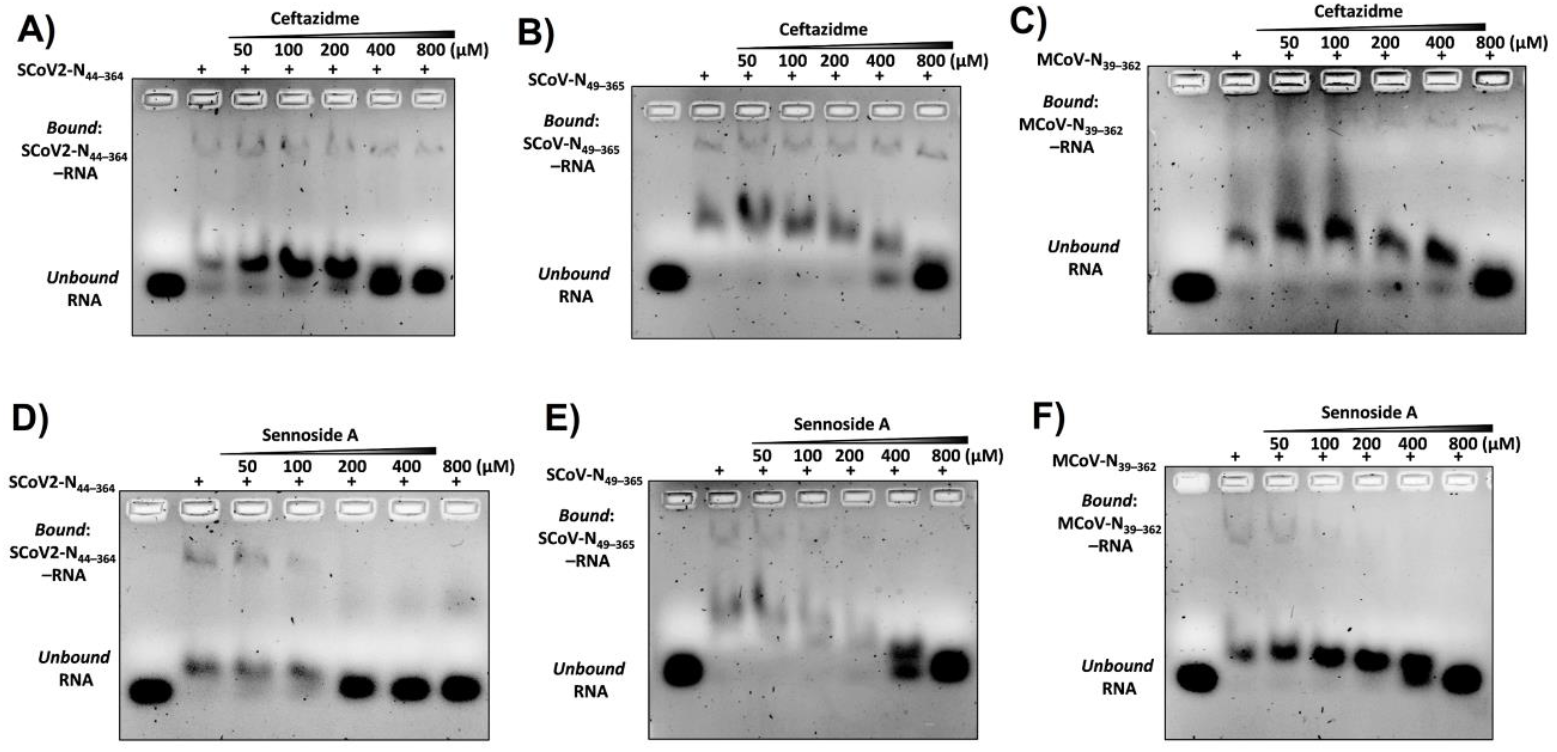
Broad-spectrum inhibition of Nucleocapsid–RNA interactions by Ceftazidime and Sennoside A across three human coronaviruses, assessed using EMSA. Preincubation of SCoV2-N_44–364_, SCoV-N_49–365_, and MCoV-N_39–362_ with increasing concentrations of Ceftazidime (A–C) or Sennoside A (D–F) resulted in dose-dependent loss of the bound complex and reappearance of free RNA, demonstrating the effective inhibition of RNA binding.

For SARS-CoV and MERS-CoV N proteins, higher concentrations (up to 800 μM) were required for full inhibition (**Fig. 4B, C**). Sennoside A showed better potency, with RNA binding inhibition at 100 μM and near-complete inhibition at 200 μM for SARS-CoV-2, and up to 400 μM for the other homologs (**Fig. 4D–F**). Notably, ceftazidime-treated samples displayed an additional band near the N–RNA complex, corresponding to a fluorescent N–ceftazidime complex, consistent with the compound’s intrinsic fluorescence upon target binding, as reported earlier [28]. These results demonstrate that both Sennoside A and Ceftazidime inhibit RNA binding across all three N protein constructs. The varying degrees of inhibitory potency across homologs may reflect subtle differences in local sequence or structural context, but the overall inhibitory action of these compounds is conserved as these compounds act through a conserved RNA-binding interface.

### 3.5. Quantitative Determination of Inhibitory Potency (IC50) for Both Compounds

Fluorescence anisotropy-based inhibition assays were performed to determine the *IC*_*50*_ of both compounds against each N protein. For ceftazidime, the *IC*_*50*_ values were 48.49 ± 2.83 μM (SARS-CoV-2), 71.13 ± 2.05 μM (SARS-CoV), and 77.96 ± 7.46 μM (MERS-CoV) (**Fig.5A–5C**). Sennoside A exhibited comparable potency, with *IC*_*50*_ values of 44.71 ± 5.34 μM, 57.95 ± 3.33 μM, and 56.58 ± 13.74 μM, respectively, for Nucleocapsid homologs (**Fig. 5E–5F**). These results have been summarized in **Table 1**. The results confirm that both inhibitors disrupt N–RNA interactions across phylogenetically distinct coronaviruses.

**Figure 5.**
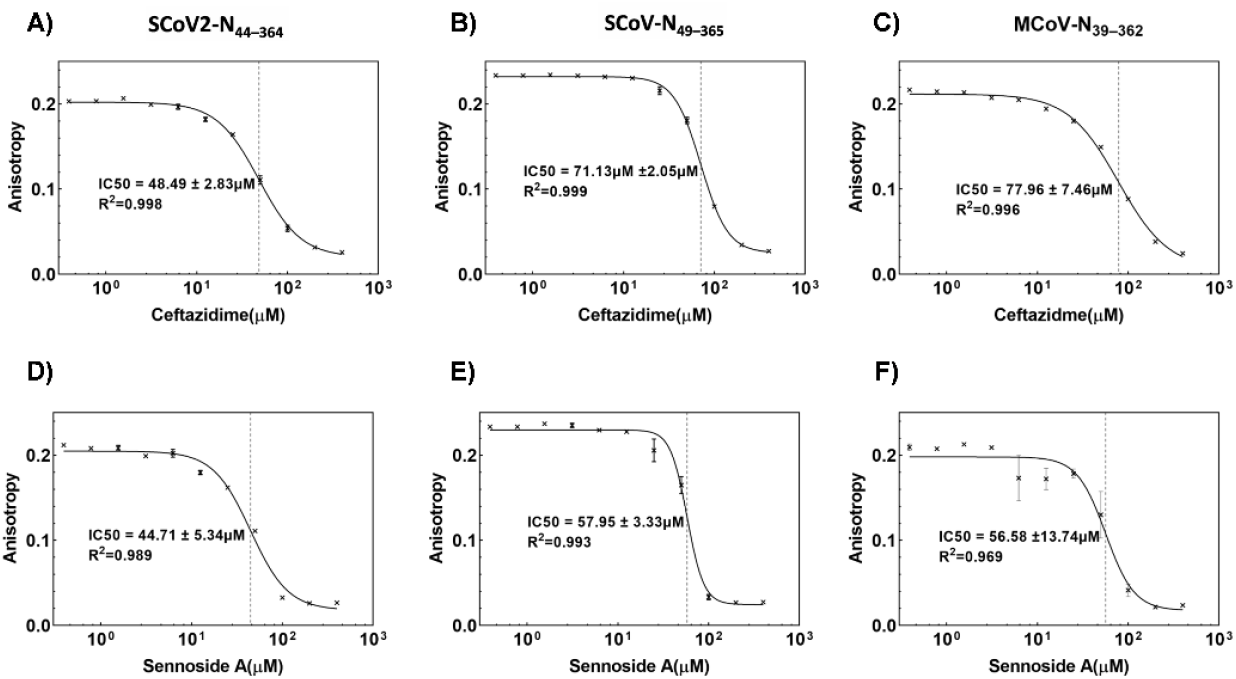
Quantitative inhibition of Nucleocapsid–RNA interactions by Ceftazidime and Sennoside. **A**. Purified nucleocapsid protein constructs SCoV2-N_44–364_, SCoV-N_49–365_, and MCoV-N_39–362_ were pre-incubated with increasing concentrations of Ceftazidime (A–C) or Sennoside A (D– F), followed by addition of a 5′-FAM-labeled RNA probe. Fluorescence anisotropy measurements showed a dose-dependent reduction in RNA binding, and sigmoidal curve fitting was used to determine the IC_50_ values of the compounds.

### 3.6. Molecular Docking Reveals Conserved Binding Modes for Both Inhibitors

To further confirm that these inhibitors act through binding to RNA binding pocket of the N-NTD, docking studies were performed using high-resolution N-NTD structures from all three viruses. Ceftazidime consistently bound to the conserved, positively charged β-hairpin loop implicated in RNA recognition. The top 200 docking poses with the MERS-CoV N-NTD yielded four distinct clusters. The most favourable cluster exhibited a HADDOCK score of −56.6 ± 1.9, with an average RMSD of 0.6 ± 0.4 Å among cluster members. The binding free energy (ΔG) of the top-ranked model was calculated to be −9.18 kcal/mol, corresponding to a predicted dissociation constant (*K*_*D*_) of ∼188 nM, and a buried surface area (BSA) of 526 Å^2^. Similarly, in the SARS-CoV N-NTD complex, Ceftazidime clustered into seven groups, with the leading cluster scoring −42.3 ± 2.4 and RMSD of 0.24 ± 0.15 Å. The calculated ΔG was −8.72 kcal/mol (*K*_*D*_ ∼416 nM; BSA: 485 Å^2^). For SARS-CoV-2, five clusters were observed, with the best cluster showing a score of −54.1 ± 2.6 and RMSD of 0.22 ± 0.12 Å. The corresponding ΔG was −8.92 kcal/mol, translating to a *K*_*D*_ of ∼296 nM, and a BSA of 452 Å^2^ (**Fig. 6A, C, E; Table 2**).

**Table 2.**
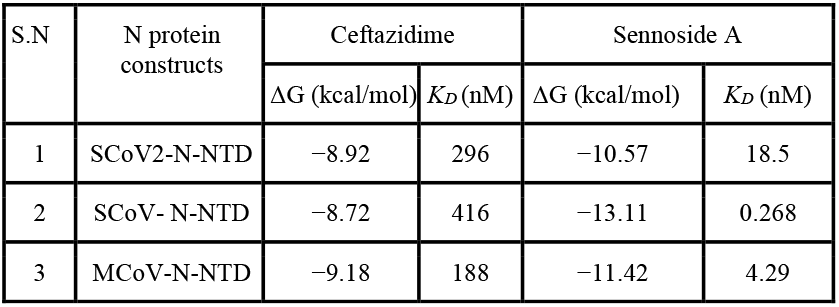
Binding free energies (ΔG) and dissociation constants (K_D_), derived from docking studies.

| S.N | N protein constructs | Ceftazidime |  | Sennoside A |  |
| --- | --- | --- | --- | --- | --- |
| | | $\Delta G$ (kcal/mol) | $K_D$ (nM) | $\Delta G$ (kcal/mol) | $K_D$ (nM) |
| 1 | SCov2-N-NTD | -8.92 | 296 | -10.57 | 18.5 |
| 2 | SCov- N-NTD | -8.72 | 416 | -13.11 | 0.268 |
| 3 | MCov-N-NTD | -9.18 | 188 | -11.42 | 4.29 |

**Figure 6.**
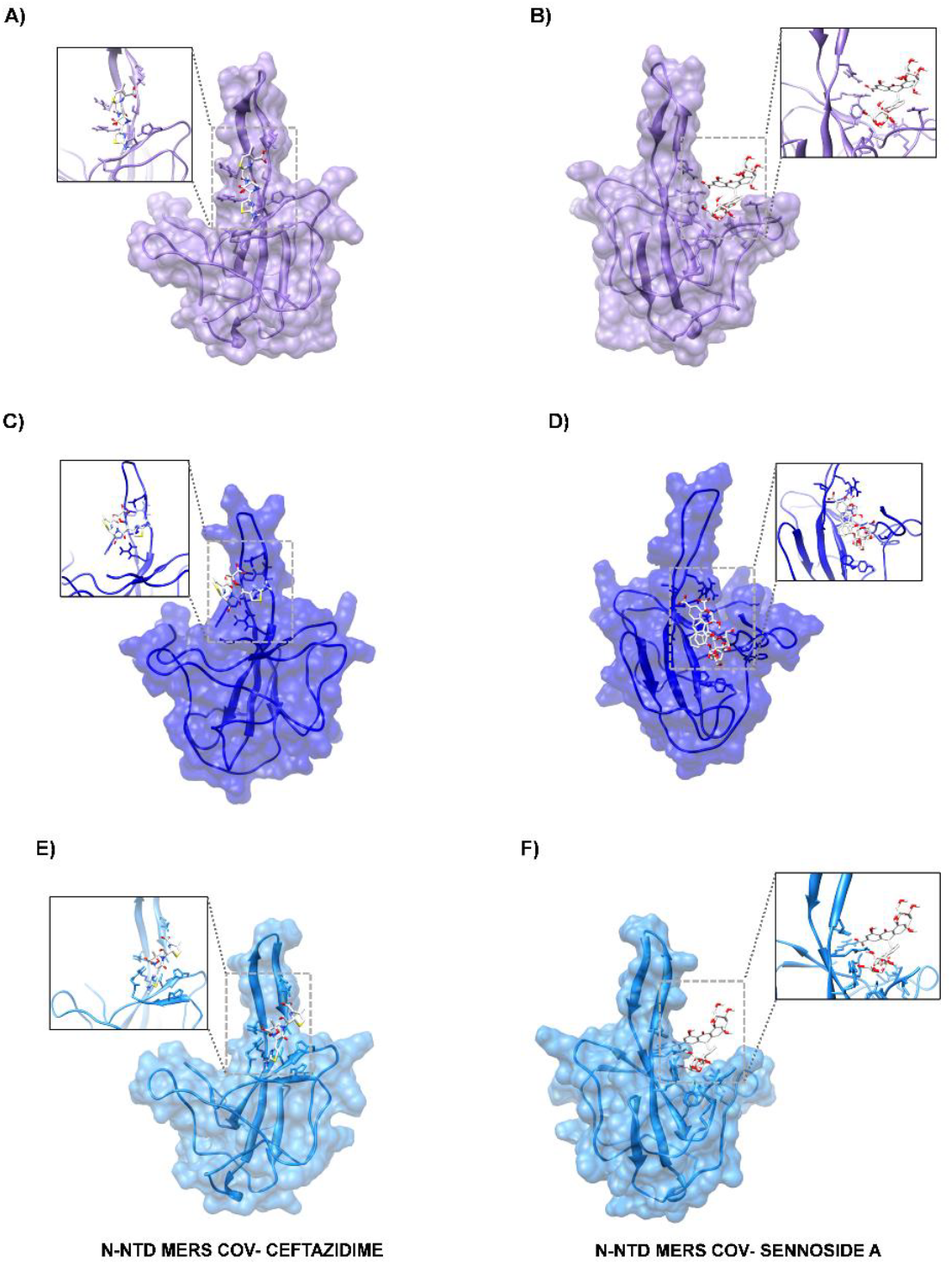
Molecular docking of Ceftazidime and Sennoside A to the N-terminal domain (N-NTD) of three human coronaviruses. (A, B) Docking poses of Ceftazidime (A) and Sennoside A (B) bound to the SARS-CoV-2 N-NTD (PDB: 8X1H). Cartoon and surface representations highlight ligand interactions with the positively charged β-hairpin “finger” and “palm” regions involved in RNA recognition. Insets show key ligand–protein interactions. (C, D) Docking of Ceftazidime (C) and Sennoside A (D) to the SARS-CoV N-NTD (PDB: 1SSK). (E, F) Docking poses of Ceftazidime (E) and Sennoside A (F) bound to the MERS-CoV N-NTD (PDB: 4UD1).

Sennoside A demonstrated stronger binding, engaging the “palm” region of the N-NTD and shielding key RNA-binding surfaces. Its predicted ΔG values ranged from −10.57 to −13.11 kcal/mol, with *K*_*D*_ values as low as 0.3 nM and larger buried surface areas compared to ceftazidime (**Fig. 6B, D, F; Table 2**). The docking results are consistent with the observed inhibitory potencies and support a conserved mechanism of action for both compounds.

## Discussion

The present study provides compelling evidence that the N-terminal domain (NTD) of the coronavirus nucleocapsid (N) protein harbors a highly conserved RNA-binding pocket that is susceptible to inhibition by small molecules. By systematically comparing the N proteins from SARS-CoV-2, SARS-CoV, and MERS-CoV, we demonstrate that both ceftazidime and sennoside A disrupt RNA binding across these phylogenetically distinct betacoronaviruses. This pan-coronavirus inhibition is underpinned by the high degree of sequence and structural conservation within the N-NTD, particularly at residues critical for RNA and inhibitor binding.

Our sequence alignment and mapping of residues reported with significant NMR chemical shift perturbation upon inhibitor binding reveal that the residues engaged by ceftazidime and sennoside A are among the most conserved within the N-NTD across SARS-CoV-2, SARS-CoV, and MERS-CoV (**Fig. 1**) [28]. Notably, these residues cluster within the basic β-hairpin “finger” and the positively charged “palm” region, both of which are essential for RNA recognition and binding. This conservation suggests that inhibitors originally identified for SARS-CoV-2 N-NTD are likely to retain efficacy against other human coronaviruses, as confirmed by our biochemical assays.

Biophysical characterization of the recombinant N protein constructs confirmed their dimeric state and solution homogeneity, supporting the validity of our functional assays. Electrophoretic mobility shift assays (EMSAs) and fluorescence anisotropy measurements established that all three N proteins bind a conserved 15-mer RNA probe with high affinity (*K*_*D*_ values ranging from ∼105 to 145 nM). Importantly, both ceftazidime and sennoside A inhibited RNA binding in a dose-dependent manner, with *IC*_*50*_ values in the low micromolar range for all three N proteins. Sennoside A consistently exhibited slightly greater potency than ceftazidime, a finding corroborated by molecular docking analyses that predicted stronger binding affinities and larger buried surface areas for sennoside A across all N-NTDs tested.

The molecular docking results further confirmed the mechanism of inhibition. Both inhibitors were found to occupy the conserved RNA-binding groove of the N-NTD, with ceftazidime primarily engaging the β-hairpin loop and sennoside A extensively shielding the palm region. These interactions likely sterically hinder RNA access to the binding site, thereby disrupting the formation of the ribonucleocapsid complex essential for viral genome packaging and replication. The convergence of inhibitor binding to these functional hotspots is consistent with our previous structural studies [28].

Targeting the N protein offers several advantages over more variable viral proteins such as the Spike glycoprotein [36]. The Spike protein is subject to rapid antigenic drift and immune escape mutations, which can compromise the efficacy of vaccines and neutralizing antibodies [17]. In contrast, the N protein is highly conserved and indispensable for multiple stages of the viral life cycle, including genome encapsidation, replication, and modulation of host immune responses [20,21,37]. The high abundance and functional indispensability of N further underscore its attractiveness as an antiviral target. Recently, several inhibitors for RNA binding of SARS-CoV-2 N protein have been reported [38,39]. Our findings align with recent SELEX and structural studies demonstrating that the N protein binds conserved RNA motifs across diverse betacoronaviruses, reinforcing the potential for broad-spectrum inhibition [10].

While our study provides strong biochemical and structural evidence for the inhibition of N–RNA interactions, it is important to acknowledge certain limitations. All assays were performed in vitro or in cell-free systems, and thus do not directly measure the impact of these inhibitors on viral replication or infectivity in cellular or animal models. Additionally, the concentrations required for inhibition are in the low micromolar range, and further optimization may be necessary to improve potency and pharmacokinetic properties for therapeutic application. Despite these limitations, our results establish a robust framework for the rational design of pan-coronavirus therapeutics targeting the N-NTD. The demonstration that small molecules can exploit conserved structural features of the N protein to achieve broad-spectrum inhibition is particularly significant in the context of ongoing and future coronavirus outbreaks. By focusing on highly conserved and functionally essential viral elements, this approach may reduce the likelihood of resistance development and provide a valuable complement to existing antiviral strategies [36,37]. The possibility of off-target effects or cytotoxicity is also limited as both the compounds are FDA approved drugs.

In summary, our study identifies the N-NTD RNA-binding groove as a conserved and druggable vulnerability in human coronaviruses. Both ceftazidime and sennoside A serve as proof-of-concept inhibitors that can disrupt N–RNA interactions across SARS-CoV-2, SARS-CoV, and MERS-CoV. Future work will focus on optimizing these scaffolds for enhanced potency and bioavailability, as well as evaluating their antiviral efficacy in cell-based and animal models. More broadly, our findings provide a template for the development of broad-spectrum antivirals targeting conserved elements of emerging and re-emerging coronaviruses, thereby strengthening global preparedness against current and future pandemics.

## Acknowledgements

We are thankful to Dr Mukesh Kumar, Head, PCS, and Dr Subhash C Bihani for insightful discussions. The work is funded by the Department of Atomic Energy, India.

## Author contribution

SS: Investigation, data analysis, manuscript writing; GDG: Conceptualization, data analysis, manuscript writing.

## Conflict of interest

None

## Notes

### Competing Interest Statement

The authors have declared no competing interest.

