## supplementary-data for "Sennoside A and Ceftazidime Inhibit Nucleocapsid RNA Binding Across SARS-CoV-2, SARS-CoV, and MERS-CoV"

***Codon optimized gene sequence of Nucleocapsid from MERS CoV & SARS CoV***

The codon-optimized gene fragments encoding the Nucleocapsid proteins from MERS-CoV (MCoV-$N_{39–362}$; residues 39–362 (UniProt ID: A0A1W5LGD9) and SARS-CoV (SCoV-N49–365; residues 49–365, UniProt ID: P59595) were chemically synthesized.

> Codon optimized gene sequence of Nucleocapsid of MERS CoV

**CATATG**AACACTGTCTCTTGGTACACTGGGCTTACCCAACACGGGAAAGTCCCTC
TTACCTTTCCACCTGGGCAGGGTGTACCTCTTAATGCCAATTCCACCCCTGCGCAAAATGCTGGGTATTGGCGG
AGACAGGACAGAAAAATTAATACCGGGAATGGAATTAAGCAACTGGCTCCCAGGTGGTACTTCTACTACACTGG
AACTGGACCCGAAGCAGCACTCCCATTCCGGGCTGTTAAGGATGGCATCGTTTGGGTCCATGAAGATGGCGCCA
CTGATGCTCCTTCAACTTTTGGGACGCGGAACCCTAACAATGATTCAGCTATTGTTACACAATTCGCGCCCGGT
ACTAAGCTTCCTAAAAACTTCCACATTGAGGGGACTGGAGGCAATAGTCAATCATCTTCAAGAGCCTCTAGCGC
AAGCAGAAACTCTTCCAGATCTAGTTCACAAGGTTCAAGATCAGGAAACTCTACCCGCGGCACTTCTCCAGGTC
CATCTGGAATCGGAGCAGTAGGAGGTGATCTACTTTACCTTGATCTTCTGAACAGACTACAAGCCCTTGAGTCT
GGCAAAGTAAAGCAATCGCAGCCAAAAGTAATCACTAAGAAAGATGCTGCTGCTGCTAAAAATAAGATGCGCCA
CAAGCGCACTTCCACCAAAAGTTTCAACATGGTGCAAGCTTTTGGTCTTCGCGGACCAGGAGACCTCCAGGGAA
ACTTTGGTGATCTTCAATTGAATAAACTCGGCACTGAGGACCCACGTTGGCCCCAAATTGCTGAGCTTGCTCCT
ACAGCCAGTGCTTTTATGGGTATGTCGCAATTTAAACTTACCCATCAGAACAATGATGATCATGGCAACCCTGT
GTACTTCCTTCGGTACAGTGGAGCCATTAAACTTGACCCAAAGAATCCCAACTACAATAAGTGGTTGGAGCTTC
TTGAGCAAAATATTGATGCCTACAAAACCTTCCCTTAA**GGATCC**

> Codon optimized gene sequence of Nucleocapsid of SARS CoV

**CATATG**AATACTGCGTCTTGGTTCACAGCTCTCACTCAGCATGGCAAGGAGGAACTTAGATTCCCTCGAGGCCA
GGGCGTTCCAATCAACACCAATAGTGGTCCAGATGACCAAATTGGCTACTACCGAAGAGCTACCCGACGAGTTC
GTGGTGGTGACGGCAAAATGAAAGAGCTCAGCCCCAGATGGTACTTCTATTACCTAGGAACTGGCCCAGAAGCT
TCACTTCCCTACGGCGCTAACAAAGAAGGCATCGTATGGGTTGCAACTGAGGGAGCCTTGAATACACCCAAAGA
CCACATTGGCACCCGCAATCCTAATAACAATGCTGCCACCGTGCTACAACTTCCTCAAGGAACAACATTGCCAA
AAGGCTTCTACGCAGAGGGAAGCAGAGGCGGCAGTCAAGCCTCTTCTCGCTCCTCATCACGTAGTCGCGGTAAT
TCAAGAAATTCAACTCCTGGCAGCAGTAGGGGAAATTCTCCTGCTCGAATGGCTAGCGGAGGTGGTGAAACTGC
CCTCGCGCTATTGCTGCTAGACAGATTGAACCAGCTTGAGAGCAAAGTTTCTGGTAAAGGCCAACAACAACAAG
GCCAAACTGTCACTAAGAAATCTGCTGCTGAGGCATCTAAAAAGCCTCGCCAAAAACGTACTGCCACAAAACAG
TACAACGTCACTCAAGCATTTGGGAGACGTGGTCCAGAACAAACCCAAGGAAATTTCGGGGACCAAGACCTAAT
CAGACAAGGAACTGATTACAAACATTGGCCGCAAATTGCACAATTTGCTCCAAGTGCCTCTGCATTCTTTGGAA
TGTCACGCATTGGCATGGAAGTCACACCTTCGGGAACATGGCTGACTTATCATGGAGCCATTAAATTGGATGAC
AAAGATCCACAATTCAAAGACAACGTCATACTGCTGAACAAGCACATTGACGCATACAAAACATTCCCATAA**GG**
**ATCC**

The restriction sites *NdeI* and *BamHI* are highlighted in bold font.

The codon-optimized gene fragments encoding MCoV-N39–362, and SCoV-N49–365 were chemically synthesized and cloned into NdeI and BamHI restriction sites of the pPCS-Trx-TEV vector, a pET-28a-derived expression vector containing an N-terminal 6xHis tag followed by a thioredoxin tag and a TEV protease cleavage site.

The plasmid **pPCS-Trx-TEV** construct sequence **(5699 bp).** The region added to pET28a vector has been highlighted in red. The NdeI and BamHI restriction sites have been underlined

TGGCGAATGGGACGCGCCCTGTAGCGGCGCATTAAGCGCGGCGGGTGTGGTGGTTACGCGCAGC
GTGACCGCTACACTTGCCAGCGCCCTAGCGCCCGCTCCTTTCGCTTTCTTCCCTTCCTTTCTCG
CCACGTTCGCCGGCTTTCCCCGTCAAGCTCTAAATCGGGGGCTCCCTTTAGGGTTCCGATTTAG
TGCTTTACGGCACCTCGACCCCAAAAAACTTGATTAGGGTGATGGTTCACGTAGTGGGCCATCG
CCCTGATAGACGGTTTTTCGCCCTTTGACGTTGGAGTCCACGTTCTTTAATAGTGGACTCTTGT
TCCAAACTGGAACAACACTCAACCCTATCTCGGTCTATTCTTTTGATTTATAAGGGATTTTGCC
GATTTCGGCCTATTGGTTAAAAAATGAGCTGATTTAACAAAAATTTAACGCGAATTTTAACAAA
ATATTAACGTTTACAATTTCAGGTGGCACTTTTCGGGGAAATGTGCGCGGAACCCCTATTTGTT
TATTTTTCTAAATACATTCAAATATGTATCCGCTCATGAATTAATTCTTAGAAAAACTCATCGA
GCATCAAATGAAACTGCAATTTATTCATATCAGGATTATCAATACCATATTTTTGAAAAAGCCG
TTTCTGTAATGAAGGAGAAAACTCACCGAGGCAGTTCCATAGGATGGCAAGATCCTGGTATCGG
TCTGCGATTCCGACTCGTCCAACATCAATACAACCTATTAATTTCCCCTCGTCAAAAATAAGGT
TATCAAGTGAGAAATCACCATGAGTGACGACTGAATCCGGTGAGAATGGCAAAAGTTTATGCAT
TTCTTTCCAGACTTGTTCAACAGGCCAGCCATTACGCTCGTCATCAAAATCACTCGCATCAACC
AAACCGTTATTCATTCGTGATTGCGCCTGAGCGAGACGAAATACGCGATCGCTGTTAAAAGGAC
AATTACAAACAGGAATCGAATGCAACCGGCGCAGGAACACTGCCAGCGCATCAACAATATTTTC
ACCTGAATCAGGATATTCTTCTAATACCTGGAATGCTGTTTTCCCGGGGATCGCAGTGGTGAGT
AACCATGCATCATCAGGAGTACGGATAAAATGCTTGATGGTCGGAAGAGGCATAAATTCCGTCA
GCCAGTTTAGTCTGACCATCTCATCTGTAACATCATTGGCAACGCTACCTTTGCCATGTTTCAG
AAACAACTCTGGCGCATCGGGCTTCCCATACAATCGATAGATTGTCGCACCTGATTGCCCGACA
TTATCGCGAGCCCATTTATACCCATATAAATCAGCATCCATGTTGGAATTTAATCGCGGCCTAG
AGCAAGACGTTTCCCGTTGAATATGGCTCATAACACCCCTTGTATTACTGTTTATGTAAGCAGA
CAGTTTTATTGTTCATGACCAAAATCCCTTAACGTGAGTTTTCGTTCCACTGAGCGTCAGACCC
CGTAGAAAAGATCAAAGGATCTTCTTGAGATCCTTTTTTTCTGCGCGTAATCTGCTGCTTGCAA
ACAAAAAAACCACCGCTACCAGCGGTGGTTTGTTTGCCGGATCAAGAGCTACCAACTCTTTTTC
CGAAGGTAACTGGCTTCAGCAGAGCGCAGATACCAAATACTGTCCTTCTAGTGTAGCCGTAGTT
AGGCCACCACTTCAAGAACTCTGTAGCACCGCCTACATACCTCGCTCTGCTAATCCTGTTACCA
GTGGCTGCTGCCAGTGGCGATAAGTCGTGTCTTACCGGGTTGGACTCAAGACGATAGTTACCGG
ATAAGGCGCAGCGGTCGGGCTGAACGGGGGGTTCGTGCACACAGCCCAGCTTGGAGCGAACGAC
CTACACCGAACTGAGATACCTACAGCGTGAGCTATGAGAAAGCGCCACGCTTCCCGAAGGGAGA
AAGGCGGACAGGTATCCGGTAAGCGGCAGGGTCGGAACAGGAGAGCGCACGAGGGAGCTTCCAG
GGGGAAACGCCTGGTATCTTTATAGTCCTGTCGGGTTTCGCCACCTCTGACTTGAGCGTCGATT
TTTGTGATGCTCGTCAGGGGGGCGGAGCCTATGGAAAAACGCCAGCAACGCGGCCTTTTTACGG
TTCCTGGCCTTTTGCTGGCCTTTTGCTCACATGTTCTTTCCTGCGTTATCCCCTGATTCTGTGG

ATAACCGTATTACCGCCTTTGAGTGAGCTGATACCGCTCGCCGCAGCCGAACGACCGAGCGCAG
CGAGTCAGTGAGCGAGGAAGCGGAAGAGCGCCTGATGCGGTATTTTCTCCTTACGCATCTGTGC
GGTATTTCACACCGCATATATGGTGCACTCTCAGTACAATCTGCTCTGATGCCGCATAGTTAAG
CCAGTATACACTCCGCTATCGCTACGTGACTGGGTCATGGCTGCGCCCCGACACCCGCCAACAC
CCGCTGACGCGCCCTGACGGGCTTGTCTGCTCCCGGCATCCGCTTACAGACAAGCTGTGACCGT
CTCCGGGAGCTGCATGTGTCAGAGGTTTTCACCGTCATCACCGAAACGCGCGAGGCAGCTGCGG
TAAAGCTCATCAGCGTGGTCGTGAAGCGATTCACAGATGTCTGCCTGTTCATCCGCGTCCAGCT
CGTTGAGTTTCTCCAGAAGCGTTAATGTCTGGCTTCTGATAAAGCGGGCCATGTTAAGGGCGGT
TTTTTCCTGTTTGGTCACTGATGCCTCCGTGTAAGGGGGATTTCTGTTCATGGGGGTAATGATA
CCGATGAAACGAGAGAGGATGCTCACGATACGGGTTACTGATGATGAACATGCCCGGTTACTGG
AACGTTGTGAGGGTAAACAACTGGCGGTATGGATGCGGCGGGACCAGAGAAAAATCACTCAGGG
TCAATGCCAGCGCTTCGTTAATACAGATGTAGGTGTTCCACAGGGTAGCCAGCAGCATCCTGCG
ATGCAGATCCGGAACATAATGGTGCAGGGCGCTGACTTCCGCGTTTCCAGACTTTACGAAACAC
GGAAACCGAAGACCATTCATGTTGTTGCTCAGGTCGCAGACGTTTTGCAGCAGCAGTCGCTTCA
CGTTCGCTCGCGTATCGGTGATTCATTCTGCTAACCAGTAAGGCAACCCCGCCAGCCTAGCCGG
GTCCTCAACGACAGGAGCACGATCATGCGCACCCGTGGGGCCGCCATGCCGGCGATAATGGCCT
GCTTCTCGCCGAAACGTTTGGTGGCGGGACCAGTGACGAAGGCTTGAGCGAGGGCGTGCAAGAT
TCCGAATACCGCAAGCGACAGGCCGATCATCGTCGCGCTCCAGCGAAAGCGGTCCTCGCCGAAA
ATGACCCAGAGCGCTGCCGGCACCTGTCCTACGAGTTGCATGATAAAGAAGACAGTCATAAGTG
CGGCGACGATAGTCATGCCCCGCGCCCACCGGAAGGAGCTGACTGGGTTGAAGGCTCTCAAGGG
CATCGGTCGAGATCCCGGTGCCTAATGAGTGAGCTAACTTACATTAATTGCGTTGCGCTCACTG
CCCGCTTTCCAGTCGGGAAACCTGTCGTGCCAGCTGCATTAATGAATCGGCCAACGCGCGGGGA
GAGGCGGTTTGCGTATTGGGCGCCAGGGTGGTTTTTCTTTTCACCAGTGAGACGGGCAACAGCT
GATTGCCCTTCACCGCCTGGCCCTGAGAGAGTTGCAGCAAGCGGTCCACGCTGGTTTGCCCCAG
CAGGCGAAAATCCTGTTTGATGGTGGTTAACGGCGGGATATAACATGAGCTGTCTTCGGTATCG
TCGTATCCCACTACCGAGATATCCGCACCAACGCGCAGCCCGGACTCGGTAATGGCGCGCATTG
CGCCCAGCGCCATCTGATCGTTGGCAACCAGCATCGCAGTGGGAACGATGCCCTCATTCAGCAT
TTGCATGGTTTGTTGAAAACCGGACATGGCACTCCAGTCGCCTTCCCGTTCCGCTATCGGCTGA
ATTTGATTGCGAGTGAGATATTTATGCCAGCCAGCCAGACGCAGACGCGCCGAGACAGAACTTA
ATGGGCCCGCTAACAGCGCGATTTGCTGGTGACCCAATGCGACCAGATGCTCCACGCCCAGTCG
CGTACCGTCTTCATGGGAGAAAATAATACTGTTGATGGGTGTCTGGTCAGAGACATCAAGAAAT
AACGCCGGAACATTAGTGCAGGCAGCTTCCACAGCAATGGCATCCTGGTCATCCAGCGGATAGT
TAATGATCAGCCCACTGACGCGTTGCGCGAGAAGATTGTGCACCGCCGCTTTACAGGCTTCGAC
GCCGCTTCGTTCTACCATCGACACCACCACGCTGGCACCCAGTTGATCGGCGCGAGATTTAATC
GCCGCGACAATTTGCGACGGCGCGTGCAGGGCCAGACTGGAGGTGGCAACGCCAATCAGCAACG
ACTGTTTGCCCGCCAGTTGTTGTGCCACGCGGTTGGGAATGTAATTCAGCTCCGCCATCGCCGC
TTCCACTTTTTCCCGCGTTTTCGCAGAAACGTGGCTGGCCTGGTTCACCACGCGGGAAACGGTC
TGATAAGAGACACCGGCATACTCTGCGACATCGTATAACGTTACTGGTTTCACATTCACCACCC
TGAATTGACTCTCTTCCGGGCGCTATCATGCCATACCGCGAAAGGTTTTGCGCCATTCGATGGT
GTCCGGGATCTCGACGCTCTCCCTTATGCGACTCCTGCATTAGGAAGCAGCCCAGTAGTAGGTT
GAGGCCGTTGAGCACCGCCGCCGCAAGGAATGGTGCATGCAAGGAGATGGCGCCCAACAGTCCC
CCGGCCACGGGGCCTGCCACCATACCCACGCCGAAACAAGCGCTCATGAGCCCGAAGTGGCGAG
CCCGATCTTCCCCATCGGTGATGTCGGCGATATAGGCGCCAGCAACCGCACCTGTGGCGCCGGT
GATGCCGGCCACGATGCGTCCGGCGTAGAGGATCGAGATCTCGATCCCGCGAAATTAATACGAC

TCACTATAGGGGAATTGTGAGCGGATAACAATTCCCCTCTAGAAATAATTTTGTTTAACTTTAA
GAAGGAGATATACCATGGGCCACCATCATCATCATCATTCTTCTGGTATGAGCGATAAAATTAT
TCACCTGACTGACGACAGTTTTGACACGGATGTACTCAAAGCGGACGGGGCGATCCTCGTCGAT
TTCTGGGCAGAGTGGTGCGGTCCGTGCAAAATGATCGCCCCGATTCTGGATGAAATCGCTGACG
AATATCAGGGCAAACTGACCGTTGCAAAACTGAACATCGATCAAAACCCTGGCACTGCGCCGAA
ATATGGCATCCGTGGTATCCCGACTCTGCTGCTGTTCAAAAACGGTGAAGTGGCGGCAACCAAA
GTGGGCGCACTGTCTAAAGGTCAGTTGAAAGAGTTCCTCGACGCTAACCTGGCCGGTACCGAGA
ACTTGTACTTCCAATCCCATATGGCTAGCATGACTGGTGGACAGCAAATGGGTCGCGGATCCGA
ATTCGAGCTCCGTCGACAAGCTTGCGGCCGCACTCGAGCACCACCACCACCACCACTGAGATCC
GGCTGCTAACAAAGCCCGAAAGGAAGCTGAGTTGGCTGCTGCCACCGCTGAGCAATAACTAGCA
TAACCCCTTGGGGCCTCTAAACGGGTCTTGAGGGGTTTTTTGCTGAAAGGAGGAACTATATCCG
GAT
